# The Ramp protocol: Uncovering individual differences in walking to an auditory beat using TeensyStep

**DOI:** 10.1101/2024.03.07.583713

**Authors:** Agnès Zagala, Nicholas E.V. Foster, Floris T. van Vugt, Fabien Dal Maso, Simone Dalla Bella

**Affiliations:** International Laboratory for Brain, Music and Sound Research (BRAMS), Montreal, Canada; Department of Psychology, University of Montreal, Montreal, Canada; Centre for Research on Brain, Language and Music (CRBLM), Montreal, Canada; School of kinesiology and physical activity sciences, University of Montreal, Montreal, Canada; Centre for Interdisciplinary Research on Brain and Learning (CIRCA), Montreal, Canada; University of Economics and Human Sciences in Warsaw, Warsaw, Poland

**Keywords:** Auditory-motor synchronization, Gait, Individual differences, Ramp protocol, TeensyStep

## Abstract

Intentionally walking to the beat of an auditory stimulus seems effortless for most humans. However, studies have revealed significant individual differences in the spontaneous tendency to synchronize. Some individuals tend to adapt their walking pace to the beat, while others show little or no adjustment. To fill this gap we introduce the Ramp protocol, which measures spontaneous adaptation to a change in an auditory rhythmic stimulus in a gait task. First, participants walk at their preferred cadence without stimulation. After several steps, a metronome is presented, timed to match the participant’s heel-strike. Then, the metronome tempo progressively departs from the participant’s cadence by either accelerating or decelerating. The implementation of the Ramp protocol required real-time detection of heel-strike and auditory stimuli aligned with participants’ preferred cadence. To achieve this, we developed the TeensyStep device, which we validated compared to a gold standard for step detection. We also demonstrated the sensitivity of the Ramp protocol to individual differences in the spontaneous response to a tempo-changing rhythmic stimulus by introducing a new measure: the Response Score. This new method and quantification of spontaneous response to rhythmic stimuli holds promise for highlighting and distinguishing different profiles of adaptation in a gait task.

## Introduction

In a public space where music is played - like a mall - you may notice that some people but not all tend to align their steps or movements to the musical beat. Spontaneous synchronization to a musical beat ^[1,2]^ is a common phenomenon that is grounded in the general tendency of auditory rhythms to engage the motor system ^[3–6]^. Humans are generally well equipped to perceive auditory rhythms, and to synchronize movements to an auditory beat. Voluntary synchronization to a beat has been thoroughly examined using finger tapping tasks, in which participants tap in synchrony with the beat of a music or a metronome ^[7–9]^. Finger tapping is the most widespread task to test voluntary synchronization, and is included in test batteries for the assessment of rhythmic abilities such as the Battery for the Assessment of Auditory Sensorimotor and Timing Abilities (BAASTA) and the Harvard Beat Assessment Test (H-BAT) ^[10–12]^. Using tapping tasks to test beat perception and voluntary synchronization is widespread and well documented ^[3,8,9,13,14]^. These studies have demonstrated significant individual differences in rhythm perception and rhythm production tasks, both with and without an external stimulus ^[12–19]^. Rhythmic abilities fall on a continuum spanning from experts, such as professional musicians, to individuals showing poor rhythmic abilities, such as beat deafness or poor synchronization ^[13,17,20–24]^. At the low end of the continuum, individuals can be impaired specifically in rhythm production ^[13]^, in rhythm perception ^[12,20]^, or both ^[17,21]^. Thus, measuring both rhythm perception and motor production is paramount to account for individual differences in rhythmic abilities ^[12,24]^.

### Spontaneous synchronization to the beat

Even though it is less documented than voluntary synchronization, there is growing evidence that people can also spontaneously (i.e., without explicit instruction) synchronize to an auditory beat when walking ^[25]^, running ^[26]^, clapping ^[27]^, or swaying their head or body ^[1,28]^. As in voluntary synchronization, significant individual differences are observed in spontaneous synchronization to a beat in healthy young adults, notably in finger tapping ^[29,30]^ and gait ^[25,31,32]^ as well as in clinical populations ^[33]^. For example, different responses are reported in patients with Parkinson’s disease when instructed to walk naturally in the presence of a rhythmic auditory stimulus ^[34,35]^. Some patients tend to adjust their gait cadence (number of steps / minute) in response to the external stimulus, while others display little or no adjustment to the external rhythm. Although individual responses to rhythmic stimuli during spontaneous walking are likely common, and may align with findings from voluntary tapping studies, there is a notable lack of targeted research protocols and comprehensive studies specifically investigating this phenomenon in healthy individuals. Gaining a better understanding of individual differences in spontaneous gait synchronization to a rhythmic stimulus can be a valuable source of information to better understand auditory-motor integration in complex, highly automatized, tasks such as gait ^[36,37]^. Moreover, this knowledge may have clinical value when considering that voluntary synchronization can be particularly difficult in clinical populations, including those who have limited cognitive resources ^[38,39]^. Knowing the factors that may foster a spontaneous motor response to a rhythmic stimulus can be particularly useful to devise interventions based on rhythmic stimulation and auditory-motor synchronization ^[40,41]^. Unfortunately, to date, there is no dedicated protocol or procedure for assessing individual differences in spontaneous auditory-motor synchronization during gait. What is needed is a technical solution that can capture an individual’s spontaneous cadence and test to what extent cadence is affected by manipulating stimulus features, such as stimulation rate.

### Using gait as a motor model of spontaneous synchronization

Gait is an excellent candidate behavior, as compared to the more widely used finger tapping ^[7,8]^, for detecting individual differences in spontaneous auditory-motor synchronization. An important drawback of finger tapping as tested in the laboratory is that this task is less ecological, a fact which limits the generalization of findings to everyday life and to more automatic rhythmic tasks. In contrast, gait is an excellent motor model protocol, as walking is an everyday, whole-body activity that emerges early during development ^[42–46]^, and remains very stable in terms of cadence and gait pattern through the life span ^[47,48]^. Gait motor control is strongly correlated with autonomy and life quality, as gait stability is a critical measure in clinical populations and the elderly ^[49–52]^. Furthermore, there is already evidence that these differences in the spontaneous gait response to a rhythmic stimulus can be detected in adults with movement disorders (for example with approximately 70% or responders and 30% of non-responders in Parkinson’s disease) ^[34,53]^.

The goal of the present study is to assess individual differences in auditory-motor synchronization in a gait task and better understand the influence of an external rhythmic stimulus on individuals’ gait cadence. To this aim we pursued two objectives. First, in order to measure individual differences in gait response to an external rhythmic stimulus, we developed and validated a new portable device (TeensyStep) that measures spontaneous cadence in real time and delivers rhythmic stimuli based on the cadence detected online. Second, we introduced and validated a new procedure (the Ramp protocol), by building on the TeensyStep technology. The protocol detects spontaneous gait responses to an auditory metronome that begins in synchrony with the participant’s spontaneous cadence and then departs from this rate progressively. As the outcome of the Ramp protocol, we proposed a measure capable of quantifying on a continuous scale the spontaneous response to a rhythmic stimulus while walking, named the “Response Score”.

## Method

### Participants

Twelve young adults (8 females), between 18 and 40 years of age (mean = 26.2 years; *SD* = 4.2) participated in the experiment. Inclusion criteria for participating in the experiment were the absence of any auditory or motor disorder that could affect walking, and normal or corrected-to-normal vision. The study was approved by the Ethics Committee of Research in Education and Psychology of the University of Montreal (CEREP-20-033-D) and carried out in accordance with the Declaration of Helsinki. Participants provided informed written consent before starting the experiment. Participants received financial compensation in exchange for their time.

### Methods for the validation of the *TeensyStep* device

We designed the TeensyStep device to 1) detect a participant’s heel strikes in real time, 2) compute the inter-step interval (i.e., the time in ms measured between one step onset and the following one), 3) deliver auditory stimuli that start at the same rate and in alignment with the participant’s initial cadence. To facilitate replication, we provide the open-source program source code along with instructions for building the full device at the GitHub URL https://github.com/dallabella-lab/teensystep, making this tool accessible to all.

TeensyStep is inspired by the TeensyTap device, which was developed for measuring finger tapping with or without external stimulation ^[54]^. TeensyStep is portable, small (4 × 4 cm) and lightweight (<75 g) and has a sampling rate of 1000 Hz (Figure 1). The detection of step events in TeensyStep is accomplished by a force sensitive resistor (FSR) positioned in the shoe under the heel of the participant’s dominant leg (Figure 1). A FSR is a sensor that changes its resistance when force is applied to its active surface. The analog force signal collected by the FSR is transmitted in real time to a Teensy 3.2 microcontroller board (PJRC, Portland, Oregon, http://www.pjrc.com/, Figure 1b) which is coupled to a compatible audio extension shield to provide auditory stimulation during the trial. When the participant steps on the FSR, the force applied to the sensor increases, resulting in a change in electrical resistance which is detected by the microcontroller and compared with a pre-set appropriate threshold of the force-resistance output to determine the step onset. This threshold has been determined prior to this study to be above the baseline output (force applied on the FSR during the dominant foot swing) for a variety of participants and as low as possible to detect the steps with the smallest delay possible (Figure 2). Before starting the testing, we checked that this threshold produced reliable step detection for each participant. As described in the Ramp protocol section, the TeensyStep device then delivered auditory stimuli synchronized to the participants’ spontaneous cadence or at various cadences according to the purpose of the study.

**Figure 1.**
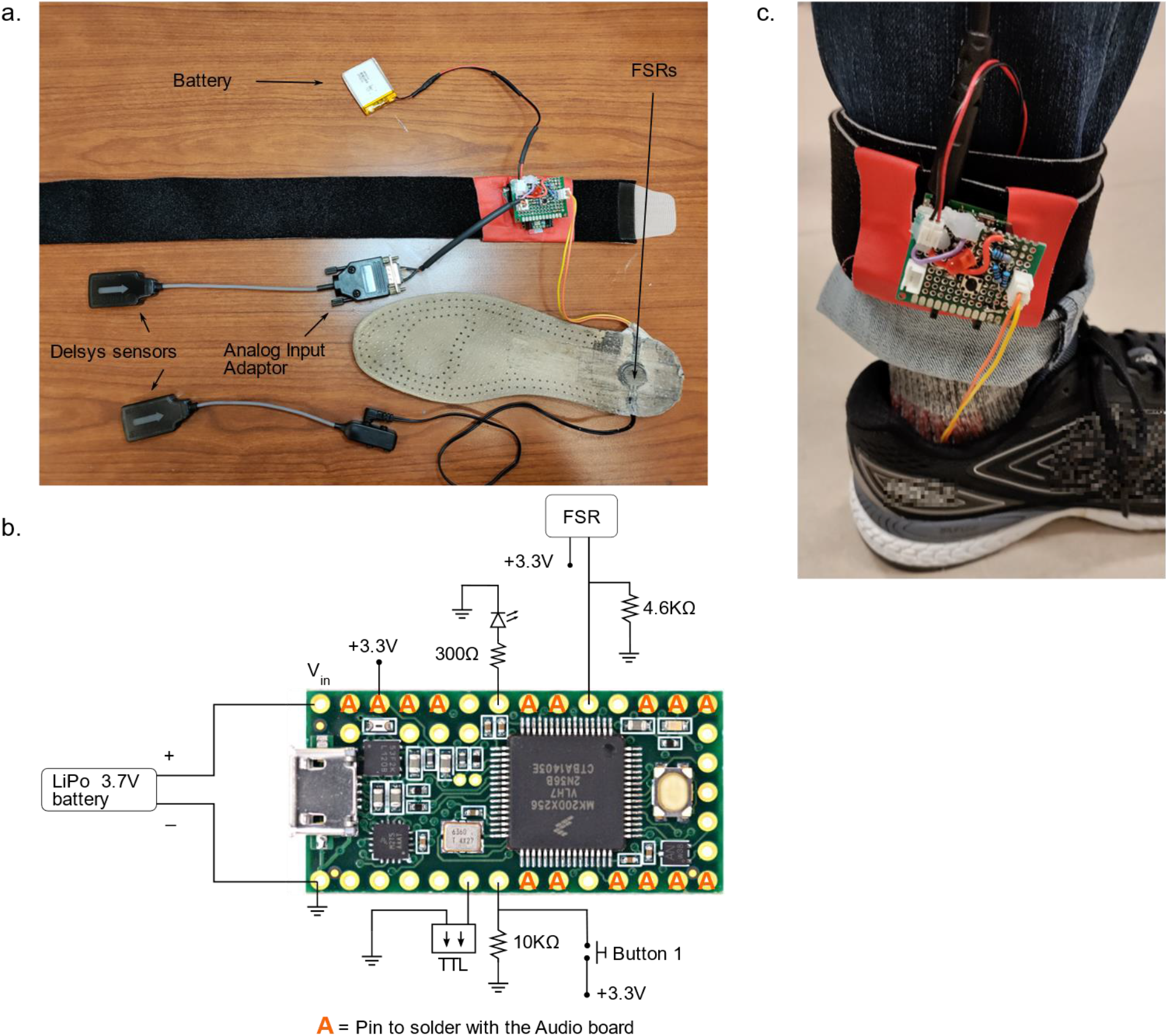
**a**. TeensyStep device (board, FSR, battery, and straps to attach the system to the participant’s ankle) and the Delsys sensors used to validate TeensyStep. **b**. Schematic of the wiring of the TeensyStep, connected to an external battery. Note that the analog input adaptor and the Delsys sensors (shown in **a**) were installed for validation purpose but are not part of the TeensyStep device. **c**. Picture of TeensyStep on a participant’s ankle.

**Figure 2.**
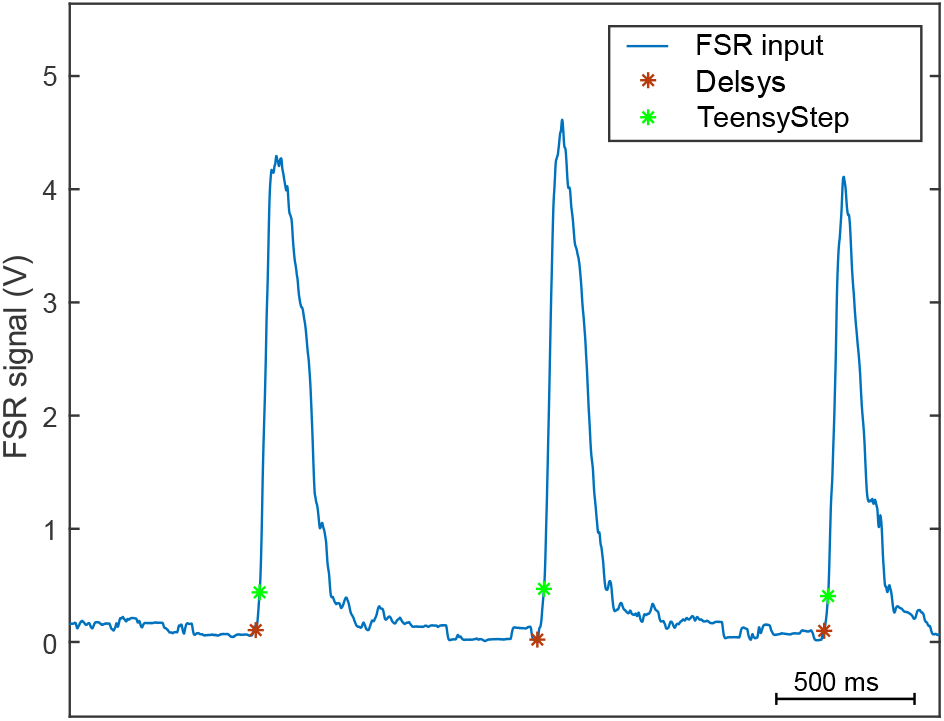
Representative time history of the Delsys FSR signal for one participant during 3 successive steps of a single trial. Each asterisk represents a step onset detection obtained from Delsys FSR and from TeensyStep.

#### Delsys FSR and step detection

To validate the accuracy of the TeensyStep device, we investigated whether the steps detected by the TeensyStep device matched those that were detected with an existing gold-standard solution. We inserted an additional FSR sensor under the heel of the participant. This FSR sensor was connected to a Delsys Trigno 4-Channel FSR Adapter, and data were recorded at a sampling rate of 1926 Hz. The Delsys system has been validated for reliability and temporal precision ^[55,56]^ and is considered a gold standard for gait analysis ^[57,58]^. The Delsys FSR allows recording of the continuous force-resistance output during gait, but it was not designed to provide real-time analysis. Accordingly, we analyzed the Delsys FSR waveforms offline to determine the step onsets. To synchronize Delsys and TeensyStep data, the TeensyStep device sent a TTL pulse from the Teensy to Delsys via an analog input adaptor that was recorded synchronously to the Delsys FSR data. To validate the TeensyStep heel strike detection procedure, we first performed offline step detection on the Delsys FSR signal waveform. For each step cycle, we detected the first point with a positive first derivative preceding the peak of the step (i.e., the first inflection onset of the Ո-curve of the step; red asterisk, Figure 2). This step detection procedure was optimized to detect step as early as possible (i.e., as close as possible to the heel strike rise of the force). This automated procedure was carried out entirely offline, after data collection was completed.

#### Validation analysis

To validate the TeensyStep device, we compared the cadence data obtained with our new device to the measurements obtained with Delsys FSR sensors. We calculated the inter-step intervals (ISIs) and the absolute timing of the step onsets throughout a trial obtained using the TeensyStep. Then, we compared these values to the ones obtained with Delsys to evaluate the precision of TeensyStep using Bland & Altman analysis ^[59–61]^. Bland & Atman analysis compares two measurement approaches, such as a gold standard and a novel method, and calculates a reproducibility coefficient (i.e., the limit of agreement between both systems) ^[59–62]^. When using Bland & Altman analysis, in order to conclude that a system is in agreement with the gold standard, 95% of the data points should be centered around the mean difference with the ground truth, and should remain within an interval of ±1.96\**SD* of this difference ^[62]^. This means the risk that a data point is not within an interval of ±1.96\**SD* around the mean difference (i.e., reproducibility coefficient in %) is lower than 5%. To compare TeensyStep to Delsys, we also calculated the percentage of the variance of one system explained by the gold standard (i.e., the coefficient of determination) to ensure that the variability of TeensyStep data is explained by the intrinsic variability of gait and not an error in our measure. However, according to Giavarina ^[62]^, the coefficient of determination between the same measure from two systems can be biased by the fact that both methods are measuring the same process or task, which could lead to an overestimation of the correlation between the two systems. Thus, it is critical to also use Bland & Altman analysis to measure the mean and the standard deviation of the difference between the gold standard and the evaluated measure to validate the agreement between methods. Consequently, we measured an average coefficient of determination (r^*2*^) between Delsys and TeensyStep ISIs as well as absolute step onset detection in order to estimate how much of the TeensyStep ISIs variance can be explained by Delsys ISI values. We also obtained a reproducibility coefficient from Bland & Altman analysis (i.e., the risk that a data point is not included within an interval of ±1.96\**SD* around the mean).

### The Ramp protocol to quantify the response to a rhythmic stimulus

As a next step, we devised a procedure, which we called “Ramp protocol”, to test whether participants walking to a rhythmic auditory stimulus (i.e., a metronome) adapt their cadence or not to a progressive tempo change in the stimulus. This protocol, implemented using the TeensyStep device, included 3 consecutive phases (Figure 3), namely, 1) estimation of a person’s initial cadence; 2) using this cadence estimate to present a steady beat sequence synchronized to the participants steps (as detailed in the *TeensyStep device* section); 3) progressive manipulation of stimulus rate (ramp). In detail: Phase 1: after the 8^th^ dominant step onset was detected, the average cadence was calculated by the microcontroller based on the 5 last steps (because the first ISIs are variable during gait initiations; ^[63]^). Phase 2: in correspondence to the 9^th^ dominant step onset, the auditory stimulus (i.e., first tone of a metronome consisting of a 100-ms woodblock sound repeated with an inter-onset interval equal to the estimated cadence) was presented. The previous estimation of the cadence allowed the TeensyStep device to start the auditory stimulus in synchrony with the participant’s spontaneous cadence in this second phase by predicting the next step onset and maintaining a tempo corresponding to the initial cadence. In parallel, the TeensyStep device recorded all step onsets, which allowed for offline analysis of stimulus tempo change on gait cadence. Phase 3 was the ramp phase, during which three experimental conditions were created by manipulating the stimulus tempo to either accelerate, decelerate, or remain constant (no change - control condition). In the ramp phase, the metronome tempo started increasing (accelerating) or decreasing (decelerating) at the 16^th^, 18^th^, or 20^th^ step of the dominant leg. The starting step number for the ramp was chosen randomly from these alternatives to avoid anticipation of the stimulus tempo change. During the ramp, the stimulus interval was increased or decreased compared to the initial cadence for each successive stimulus by a duration corresponding to 0.5% of the baseline inter-stride interval. The ramp continued for 40 stimuli, reaching a stimulus interval that was 20% shorter or longer (faster or slower rate) relative to the participant’s preferred cadence measured during the first phase. We considered the no-change condition as a control condition in which the stimulus tempo remained at the participants’ initial cadence during the third phase. For each of these ramp conditions (acceleration, deceleration, and control), participants were asked to either synchronize to the stimulus (synchronization block) or walk naturally with the stimulus (natural block). The natural instruction was used to evaluate the spontaneous response of the participant to a change of the stimulus, and the synchronization instruction evaluated whether the participant was able to synchronize to the stimulus. Each block included 10 trials: 4 with acceleration, 4 with deceleration, and 2 control trials with no tempo change. Because the natural instruction is the most sensitive to bias depending on which instructions came first, all participants started the experiment with the natural block. Trial order (i.e., acceleration, deceleration and no change) within a block was randomized. For the purpose of the present study, only 2 experimental conditions (acceleration and deceleration during the natural block) were used for the validation analysis. The results from the synchronization block were used to ensure that participants could respond to the stimulus change if instructed. Each tempo change was repeated 4 times in a random order.

**Figure 3.**
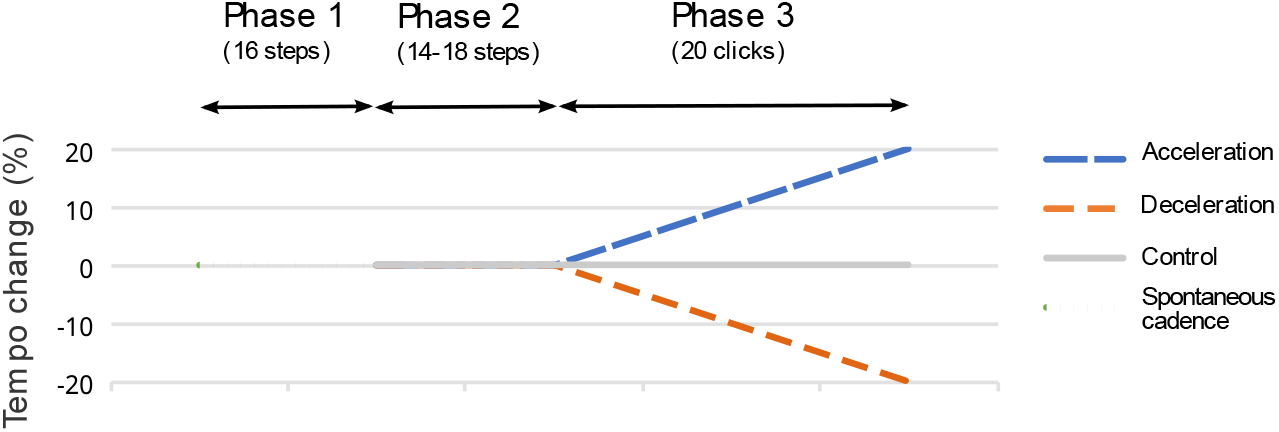
Sequence of a trial depending on the tempo change: acceleration or deceleration up to 20% of the spontaneous cadence, or no change. Tempo change is indicated in percent difference in stimulus inter-onset interval compared to the phase 1 baseline, where positive values represent tempo increase (shorter intervals). Phase 1 corresponds to the spontaneous cadence with no rhythmic cue, phase 2 begins when the audio stimulus starts (synchronized to the initial cadence), and phase 3 begins with the tempo change (acceleration, deceleration, or no change).

#### Measure of the individual response to the stimulus tempo change

In order to quantify the individual response while walking to a rhythmic stimulus with a tempo change (acceleration or deceleration of the metronome), we analyzed the ISI change values measured at each stride heel strike in the spontaneous instruction condition. The ISI change is expressed as a percent relative to the average ISI (ISI_0_) measured in the first phase of the trial, i.e., 100*(ISI_i_ – ISI_0_)/ISI_0_. These ISI change values were then used in regression models (with time in seconds as x, using the “fitlm” function in Matlab) and area calculations. The steps needed to obtain a measure of the individual response (so called “Response Score”) are illustrated in Figure 4a. First, a precondition for including a trial in the calculation was a non-significant regression slope for ISI change vs. time during the second phase (walking with the metronome at constant initial tempo); i.e., the participant’s cadence should not have started changing prior to the ramp. The ISI change values in the third phase (the ramp) were averaged across trials at each heel strike and used to calculate the Response Score. This score corresponded to the area between the acceleration and deceleration curves of ISI change across heel strikes in the third phase, divided by the area between the metronome curves (Figure 4b). An over-correction (i.e., accelerating or decelerating even more than the stimulus) would lead to a percentage exceeding 100%. On the contrary, negative scores could be obtained when participants tended to compensate the change in the opposite direction to the change (i.e. accelerating when the metronome is decelerating and/or vice versa).

**Figure 4.**
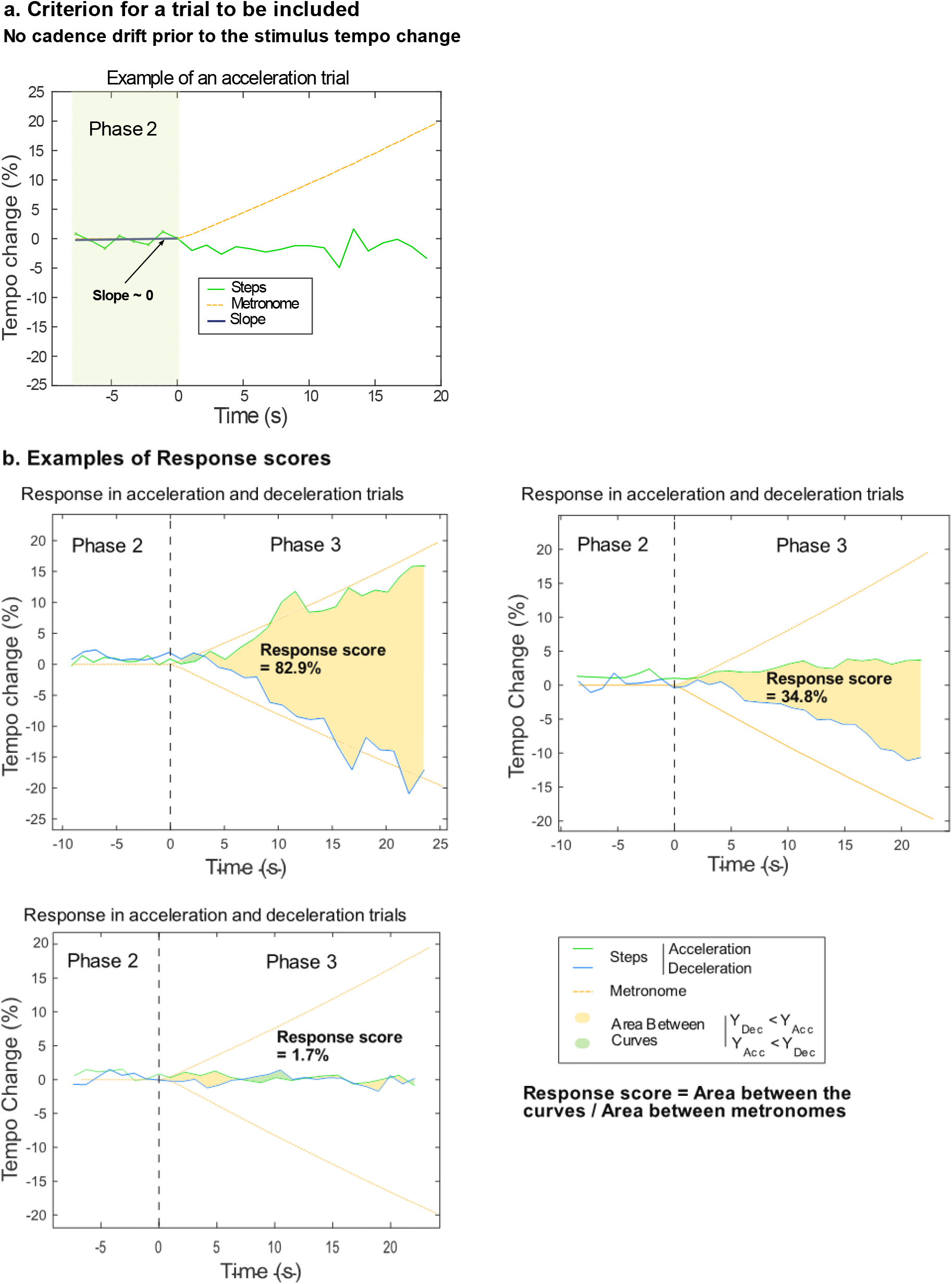
a. Criteria to ensure the specificity of a change in the cadence in order to calculate the Response Score. b. Response Score for 3 participants displaying various responses. In a. and b., the time 0 corresponds to the beginning of the third phase (i.e., the ramp phase) in order to highlight the response of the participant to the tempo change of the metronome.

Finally, we validated the robustness of the Response Score, by ensuring that the score cross-validates across trials. For each participant we computed the score multiple times using a leave-one-out procedure. Each time the calculation was performed without including a single trial, a procedure repeated for all the trials. Given that each participant produced at least 2 trials per tempo condition (acceleration and deceleration), we calculated the Response Score N_tot_ times (N_tot_ = N_acc_ + N_dec_) per participant. This procedure allowed us to assess the contribution of each trial individually and to observe the consistency of the response quantification dependent on the individual trial contribution. We compared the Response Score for each participants obtained using all the trials and the mean Response Score obtained using the leave-one-out procedure.

## Results

### Validation of TeensyStep device

#### Relative Timing – Inter-Step Interval comparison

In order to validate TeensyStep, we compared the ISIs obtained via TeensyStep and Delsys FSR. Using data from the whole sample, we obtained *r*^*2*^ = .99 (Figure 5, left) and reproducibility coefficient = 1.8% (Figure 5, right). The limit of agreement, i.e., ±1.96\**SD*, is ±19.4 ms and the mean difference is not significantly different from 0 (*t*(9390) = 0.23; *p* = .82). Using individual data, we obtained an average *r*^*2*^ = .95 (*SD* = 0.03; range = .89-.99). The average reproducibility coefficient in our study is 1.8% (*SD* = 0.67; range = 0.83%-3.20%). The individual data for each participant are presented in Supplementary materials (Appendix SA1 and SA2).

**Figure 5.**
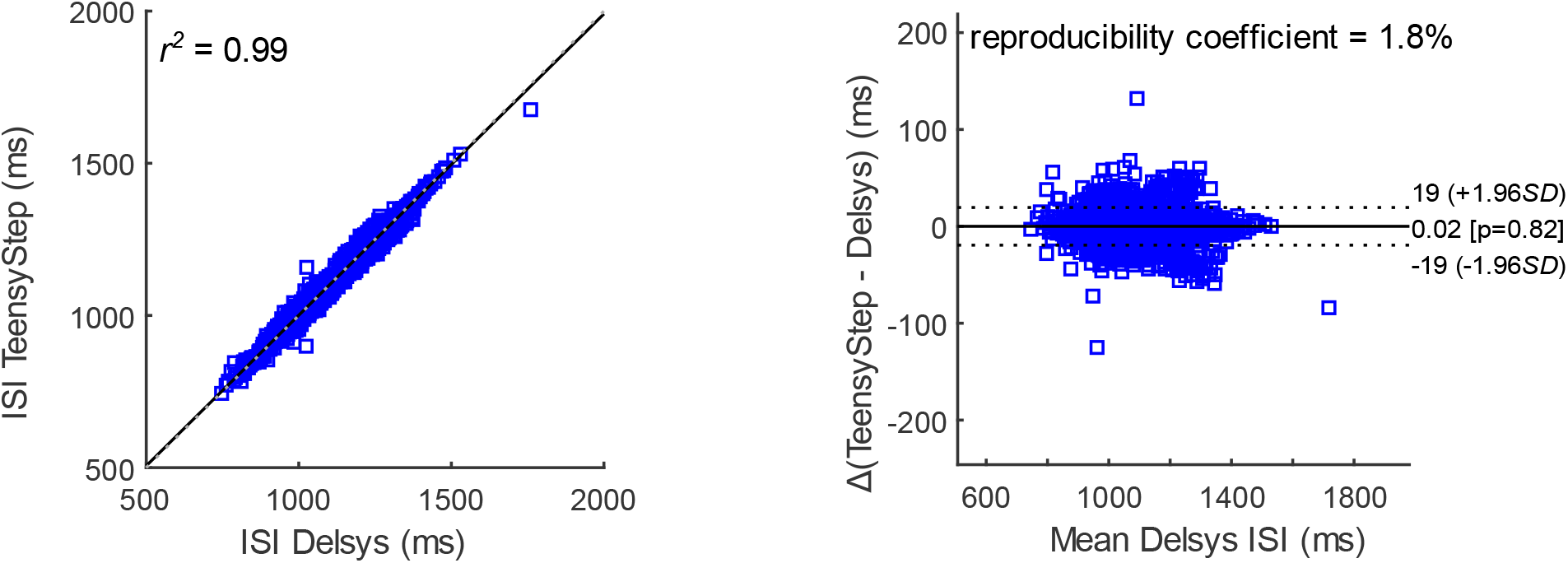
Similarity (left) and Bland & Altman representation (right) of the ISIs measured using the Delsys and TeensyStep. On the left panel, the slope of the regression is y = 0.99x+11.1. The x axis of the right panel represents the absolute delay in ms between the onsets calculated by both TeensyStep and Delsys.

#### Absolute Timing – Step onsets comparison

We also compared the step onsets detected via TeensyStep FSR and Delsys FSR. Using data from the whole sample, we obtained *r*^*2*^ = 1.0 and reproducibility coefficient = 0.21% (Figure 6). The limits of agreement, i.e., ±1.96*SD, are respectively +40.9 and -35.7 ms and the mean difference is 2.6 ms (*t*(9701) = 13.10; *p* < .001). When doing this analysis on individual data, we obtained an average *r*^*2*^ = 1.0 (*SD* = 0.0). The average reproducibility coefficient in our sample is 0.21% (*SD* = 0.09; range = 0.07%-0.33%). The average of the mean difference between both systems in term of absolute timing is 7.9 ms (*SD* = 4.3; range = 1.2-14 ms). The individual data for each participant are presented in Supplementary materials (Appendix SA3).

**Figure 6.**
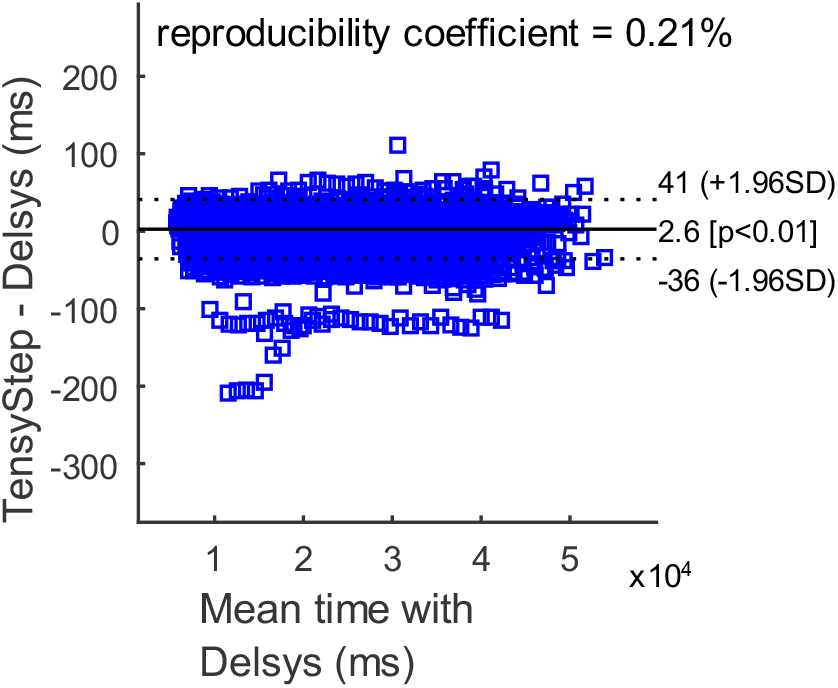
Comparison of the absolute step onsets data of the participants obtained with both FSRs from TeensyStep and Delsys (used as a gold standard). This represents the limits of agreement between both systems. These are the data for the 12 participants.

### Validation of the Ramp protocol and of the Response Score

We tested whether the Ramp protocol can provide a measure (i.e., the Response Score) capable of capturing the spontaneous response to a rhythmic stimulus while walking. We applied the response criteria described in Figure 4a to the 8 trials (4 per condition) for each of the 12 participants. The exclusion criterion for a significant change slope in phase 2 (Figure 4a) resulted in discarding either 0 trials (*n* = 4 participants), 1 trial (*n* = 6) or 2 trials (*n* = 2) per participant. The mean Response Score equals 15.26% (range: -14.4% to +82.9%; *SD* = 27.03) (Figure 7). As can be seen in Figure 7, there seems to be a continuum in the Response Score, with 8/12 of the participants around 0% (between -20% and + 20%), and 4/12 above +20%, pointing to a spontaneous response to the rhythmic stimulus. Even though the sample is quite small, we observe one Response Score above 80 indicating a strong spontaneous response and a few small negative responses. Altogether, these findings show that the Response Score is capable of capturing the range of spontaneous responses to a rhythmic stimulus, spanning from an absence of response to strong responses to the stimulation.

**Figure 7.**
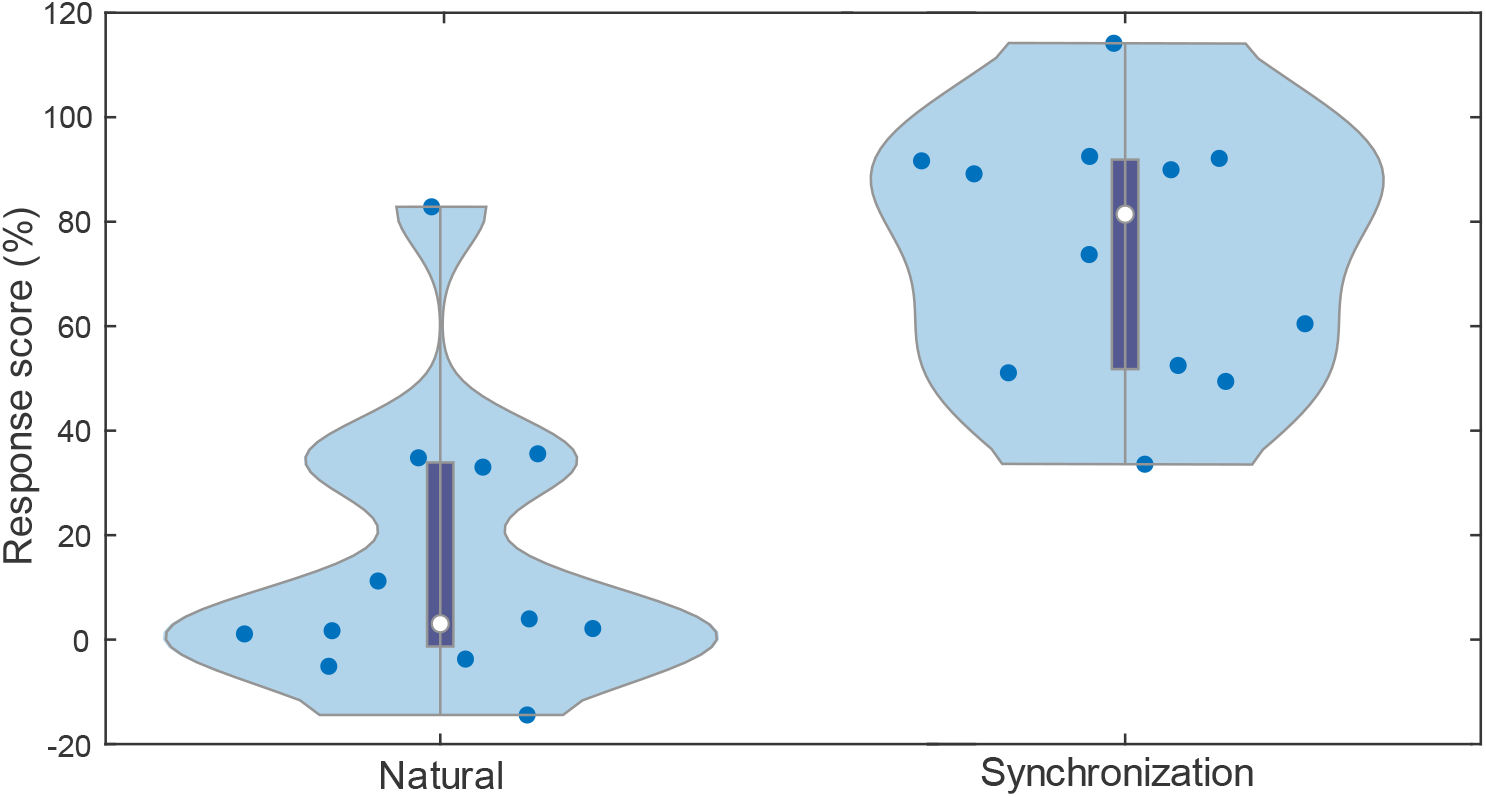
Distribution of the Response Score among the 12 participants during both Natural (i.e. no synchronization instruction) and Synchronization instructions.

In order to verify that participants could respond to the change of stimulus tempo when asked to synchronize with the stimulus, we also calculated the Response Score obtained during the synchronization block. We found that most participants could voluntarily vary their cadence to follow an external rhythmic stimulus: mean(Response Score_synchronization_) = 74.19% (range: 33.6% to 114.2%; *SD* = 24.29%). This control condition allowed us to distinguish the ability to spontaneously respond to a change in the stimulus tempo from a general inability to synchronize with a changing stimulus.

Finally, we measured the robustness of the Response Score by computing the score multiple times while leaving out a single trial (leave-one-out method). When applying this procedure on the Response Score among the 12 participants, the mean difference between these Response Scores was Δ_leave-one-out_ = -0.48% (*SD* = 0.74%). We also calculated the standard deviation of the iterations of the leave-one-out analysis for each participant and the mean deviation was mean (Standard deviation) = 3.14% (*SD* = 2.60%). Both the difference in the Response Score and its *SD* were very low thus showing that the Response Score was very stable and did not depend on a single trial.

## Discussion

The goal of the present study was to develop a method for assessing individual differences in spontaneous auditory-motor synchronization to a rhythmic stimulus, using a gait task. This method would serve to gain a better understanding of the mechanisms and factors underlying spontaneous synchronization to the beat. A first step was to validate the TeensyStep device against a ground truth, showing that the method can reliably measure gait cadence and adapt the stimulation to a participant’s natural cadence. A second step was to introduce the Ramp protocol, as a way to assess spontaneous response to a rhythmic stimulus during gait. Here we provided first evidence in a small sample of participants that the Ramp protocol can capture individual differences in the spontaneous response to a rhythmic stimulus and that this can be reliably quantified via a simple score (i.e., the Response Score).

In order to implement the Ramp protocol, which requires real-time estimation of gait cadence, we adapted the TeensyTap ^[54]^ method, initially used for finger tapping, to gait assessment. To this aim, we used a Teensy microcontroller coupled to a FSR, because it is portable, powerful for its size, and allows the collection of reliable raw data based on the FSR input and internal clock. Additionally, the use of the Teensy Arduino device provides compatibility with many accessories such as LED, response button, and high-quality audio output for the stimulus, which results in a versatile tool for a total cost as low as 100 USD. To test the temporal precision of the raw data of a FSR collected with TeensyStep, we compared data collected with our device with a validated non-real-time device, the Delsys FSR. The comparison led to a coefficient of determination higher than 0.99, and a reproducibility coefficient lower than 2%. High reliability is afforded when those criteria are above 0.95 and below 5% respectively ^[62]^. The limits of agreement were ±19 ms and -35.7 to +40.9 ms for ISI and total time, respectively. Considering that the average ISI is 1200 ms and the total time is around 45000 ms, the TeensyStep device presents high accuracy for monitoring in real time heel strike events. Thus, both Delsys and TeensyStep FSR provide very similar results in terms of period (time interval between steps) as well as phase (absolute time of the step onset). Accordingly, both the high correlation between Delsys and Teensy FSR (> 99.9%) and the low reproducibility coefficient (< 0.25%), indicate a minimal discrepancy between both methods when measuring step time. These findings are strong indications that TeensyStep can be used to assess gait cadence in real time with a very high reliability. We found no difference when comparing the ISIs from both systems.

However, we noticed a constant delay (7.9 ms) in the step detection absolute time between the systems. To our knowledge, there is no clear threshold for detecting a delay between a step and an auditory stimulus. According to Fujisaki & Nishida ^[64]^, auditory-tactile threshold detection is approximately 10 Hz for isochronous pulses in both modalities and 30 ms for single pulse (i.e., a single pitch change in a continuous sound). Another study in auditory-tactile perception of synchrony shown that the temporal threshold to perceive a keystroke and a following tone as simultaneous is 102±65 ms for musicians, and 180±104 ms for non-musicians ^[65]^. Then, a delay of 7.9 ms should not be perceptible. In sum, our system is as temporally precise as Delsys step onset detection FSR, while TeensyStep presents the advantage of being usable in real-time. Despite significant inter-individual differences in terms of spontaneous cadence and the response to the stimulation, this study showed that the results from TeensyStep are also very robust.

In the Ramp protocol we tested the adaptation of the gait cadence to a change of an external rhythmic stimulus. The protocol was individualized, as the stimuli (metronome clicks) were presented relative to the participant’s preferred cadence, a process afforded by real-time cadence detection via TeensyStep. Adaptation of stimulation to individual participants is important, as spontaneous cadence can vary significantly among adults ^[66]^. We introduced the Response Score, a measure in percent reflecting the magnitude of the spontaneous response to a change in the rhythmic stimulus (acceleration and deceleration). In this study, the Response Score was very stable across trials, revealing very little variation in the cross-validation analysis. Among this small sample of participants (n = 12) we showed that the Response Score is sensitive to the entire range of responses with a high inter-individual variability. Out of the 12 tested participants, 8 showed a Response Score around 0% (range: -14.4% to 11.2%), indicating a lack or very weak response to the stimulus change; while the other 4 showed a Response Score greater than 25% (range: 33.0% to 82.9%), including one participant with a Response Score above 80%, showing a strong spontaneous response to the stimulus change. However, a Response Score of 100% does not always mean perfect synchrony since the Response Score expresses the magnitude if the change in cadence relative to the external tempo change rather than a measure of absolute synchronization. Among young adults, the fact that gait is so automatized could partly explain why most participants show little spontaneous response to the stimulus change, even though they are capable of synchronizing to the stimulus, if instructed. Gait, one of the most practiced and important movements allowing one to move independently and autonomously ^[50,51,67]^, is highly automatized and energetically cost-efficient, especially when the walking pace is within the range of our individual preferred cadence ^[68–71]^. In spite of its stability, we see significant inter-individual differences in the way healthy young adults adapt their cadence in response to a rhythmic auditory stimulus, no matter the instruction ^[32,72]^. In our sample, some of the participants showed a slightly negative Response Score. Negative Response Scores, while they may appear paradoxical, may reflect a small degree of noise in the measure, confirming a lack of response or a tendency to compensate the tempo change by spontaneously changing cadence in the opposite direction of the stimulus change. In summary, the findings from the Ramp protocol are highly promising, revealing the potential of capturing individual variations in spontaneous synchronization to a beat through the Response Score.

If this sensitivity of the Response Score to individual differences is confirmed with a larger sample, it may enable the identification of participants or patients who are especially responsive and can be chosen as candidates for rhythmic interventions, particularly in tasks related to gait. Recent studies show that auditory-motor synchronization can have a positive impact on patient’s gait, even on a relatively long term ^[73,74]^, but only in approximately 50% of patients with a stimulus tempo close to their spontaneous cadence ^[34,53,35]^. In order to understand these individual differences and to predict whether a person can benefit from auditory rhythmic stimulation while walking, we need to take into account the spontaneous cadence of participants (i.e., which tempo is comfortable for a given individual) and their spontaneous tendency to respond to an external pace (i.e., depart from their spontaneous cadence). Using a metronome that gradually departs from the spontaneous cadence allows us to evaluate a window of stability around the preferred cadence, i.e., how much can a participant depart from this initial pace under conditions where the instruction is to synchronize or to walk naturally. The main advantage of TeensyStep is that the assessment of the cadence and the subsequent synchronization of the metronome is performed in real time, which is a very ecological task for the participant or the patient.

In spite of these promising results, the are some limitations for the method that should be taken into account in future studies. Some limitations concern the TeensyStep method. A condition for achieving accurate real-time alignment of audio stimuli with the participant’s heel strike is that initial step intervals are regular, thus ensuring reliable prediction. Secondly, the FSR threshold determined for step detection may vary as a function of the specific experimental setting, the type of shoes or insoles and the pattern of gait cycle (e.g. heel strike to forefoot spectrum). Before running a new experiment is thus advised to check the FSR waveform to make sure the threshold provides a reliable measure considering the new setting, or if it needs to be modified. Incorrect thresholds may delay detection, impacting step onset detection and cadence assessment. Finally, it is advised to consider using a Teensy 4.0 board for longer trials or for collecting numerous heel strike events, as the current RAM usage may limit data collection beyond 1 minute.

In the future, the method that we developed can be instrumental for advancing research on individual differences in a gait task. Providing an ecological and temporally precise tool to understand the conditions leading individuals to move away from their stable spontaneous cadence rather than maintaining the initial pace can shed light on important aspects of auditory-motor synchronization during gait, and the underlying constraints and mechanisms. First, it will help to quantify and categorize the different spontaneous response types, and measure how different conditions (e.g., tempo change or instructions) affect the stability of gait depending on an individual profile (e.g., initial gait variability and pace, rhythmic abilities, cognitive factors). Secondly, it may help in circumscribing the limits of this gait stability — by clarifying the circumstances (e.g., cognitive abilities, motor stability, rhythmic abilities) leading to a potential dual-task costs for gait stability. Finally, by uncovering cognitive or motor measures capable of predicting the individual response to external rhythmic stimulus, it will become possible to adapt existing rhythmic interventions for targeted clinical populations. We could notably predict the best individualized stimulation parameters to improve the stability of gait.

The results obtained in our pilot study aiming at validating the Ramp protocol should be replicated and extended with a future study with a larger sample. This will allow more accurate and reliable characterization of the distribution of the Response Score and investigate whether we can distinguish categories such as responders and non-responders based on their Response Score.

## Acknowledgment

This study was funded by a Discovery Grant (RGPIN-2019-05453) from the Natural Sciences and Engineering Research Council of Canada (NSERC), and by the Canada Research Chair program (CRC in Music Auditory-motor Skill Learning and New Technologies) to SDB, and by scholarships from the Center for Interdisciplinary Research on Brain and Learning (CIRCA) and the Center for Research on Brain, Language and Music (CRBLM) to AZ.

We wish to thank Alex Nieva for his valuable assistance in developing TeensyStep, and Florence Landry-Lehoux and Véronique Renaud for their help in participant recruitment and data collection.

## Data availability

To facilitate replication, we provide the open-source program source code along with instructions for building the full device at the GitHub URL https://github.com/dallabella-lab/teensystep, making this tool accessible to all. The datasets generated during and analyzed in this study are available from the corresponding author on reasonable request. The experiment was not preregistered.

## Author contributions

Conception of the study: AZ, SDB, FDM: writing of a first draft of the manuscript: AZ, SDB; recruitment and data collection: AZ; analysis design: AZ, NF, FVV, FDM, SDB; data analysis: AZ, FDM, NF, SDB; all authors contributed to edit the final version of the manuscript.

