## Supplementary material for "The Ramp protocol: Uncovering individual differences in walking to an auditory beat using TeensyStep"

**Supplementary material - Appendices**

***Relative Timing – Inter-Step-Interval comparison***

Individual results of the Bland and Altman analysis of the similarity (SA1) and Bland & Altman representation of the inter-step intervals measured using the Delsys and TeensyStep (SA2).

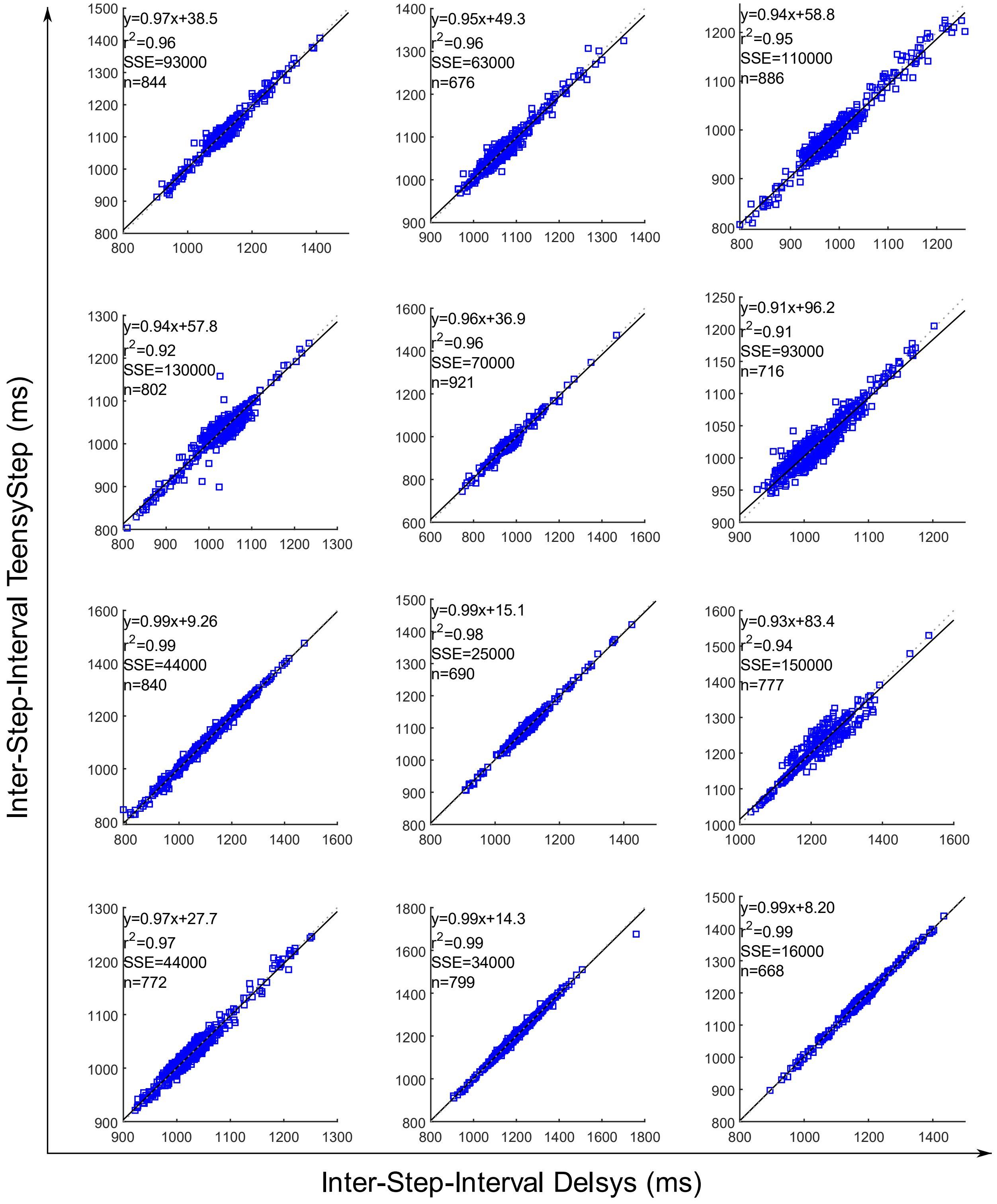

Inter-Step-Interval TeensyStep (ms)
Inter-Step-Interval Delsys (ms)
y=0.97x+38.5
r^2=0.96
SSE=93000
n=844
y=0.95x+49.3
r^2=0.96
SSE=63000
n=676
y=0.94x+58.8
r^2=0.95
SSE=110000
n=886
y=0.94x+57.8
r^2=0.92
SSE=130000
n=802
y=0.96x+36.9
r^2=0.96
SSE=70000
n=921
y=0.91x+96.2
r^2=0.91
SSE=93000
n=716
y=0.99x+9.26
r^2=0.99
SSE=44000
n=840
y=0.99x+15.1
r^2=0.98
SSE=25000
n=690
y=0.93x+83.4
r^2=0.94
SSE=150000
n=777
y=0.97x+27.7
r^2=0.97
SSE=44000
n=772
y=0.99x+14.3
r^2=0.99
SSE=34000
n=799
y=0.99x+8.20
r^2=0.99
SSE=16000
n=668

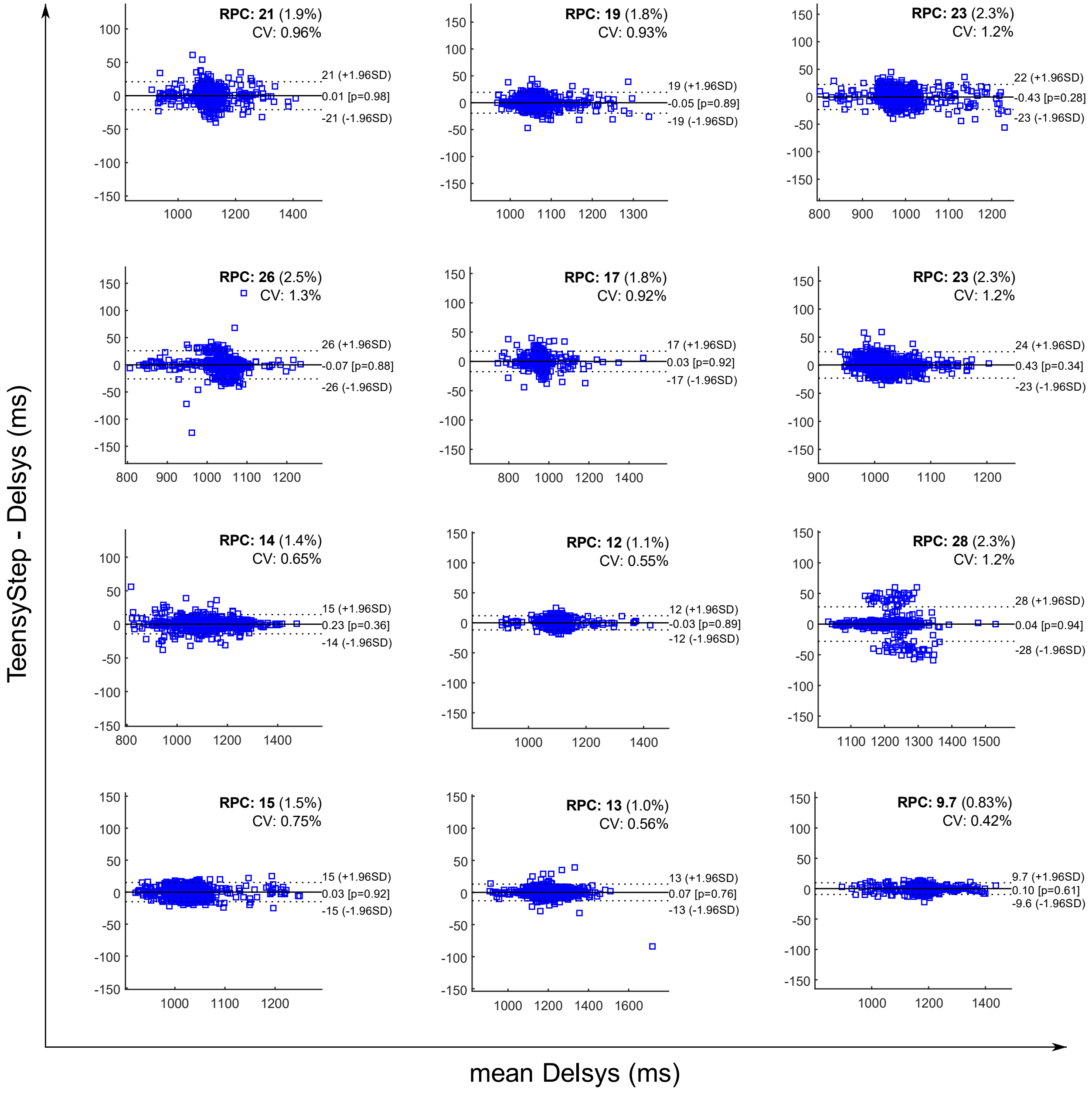

TeensyStep - Delsys (ms)
mean Delsys (ms)
RPC: 21 (1.9%)
CV: 0.96%
21 (+1.96SD)
0.01 [p=0.98]
-21 (-1.96SD)
RPC: 19 (1.8%)
CV: 0.93%
19 (+1.96SD)
-0.05 [p=0.89]
-19 (-1.96SD)
RPC: 23 (2.3%)
CV: 1.2%
22 (+1.96SD)
-0.43 [p=0.28]
-23 (-1.96SD)
RPC: 26 (2.5%)
CV: 1.3%
26 (+1.96SD)
-0.07 [p=0.88]
-26 (-1.96SD)
RPC: 17 (1.8%)
CV: 0.92%
17 (+1.96SD)
0.03 [p=0.92]
-17 (-1.96SD)
RPC: 23 (2.3%)
CV: 1.2%
24 (+1.96SD)
0.43 [p=0.34]
-23 (-1.96SD)
RPC: 14 (1.4%)
CV: 0.65%
15 (+1.96SD)
0.23 [p=0.36]
-14 (-1.96SD)
RPC: 12 (1.1%)
CV: 0.55%
12 (+1.96SD)
-0.03 [p=0.89]
-12 (-1.96SD)
RPC: 28 (2.3%)
CV: 1.2%
28 (+1.96SD)
0.04 [p=0.94]
-28 (-1.96SD)
RPC: 15 (1.5%)
CV: 0.75%
15 (+1.96SD)
0.03 [p=0.92]
-15 (-1.96SD)
RPC: 13 (1.0%)
CV: 0.56%
13 (+1.96SD)
0.07 [p=0.76]
-13 (-1.96SD)
RPC: 9.7 (0.83%)
CV: 0.42%
9.7 (+1.96SD)
0.10 [p=0.61]
-9.6 (-1.96SD)

***Absolute Timing – Step onset comparison***

Individual results of the Bland & Altman representation of the inter-step intervals measured using the Delsys and TeensyStep.

*Supplementary Appendix 3 (SA3)*

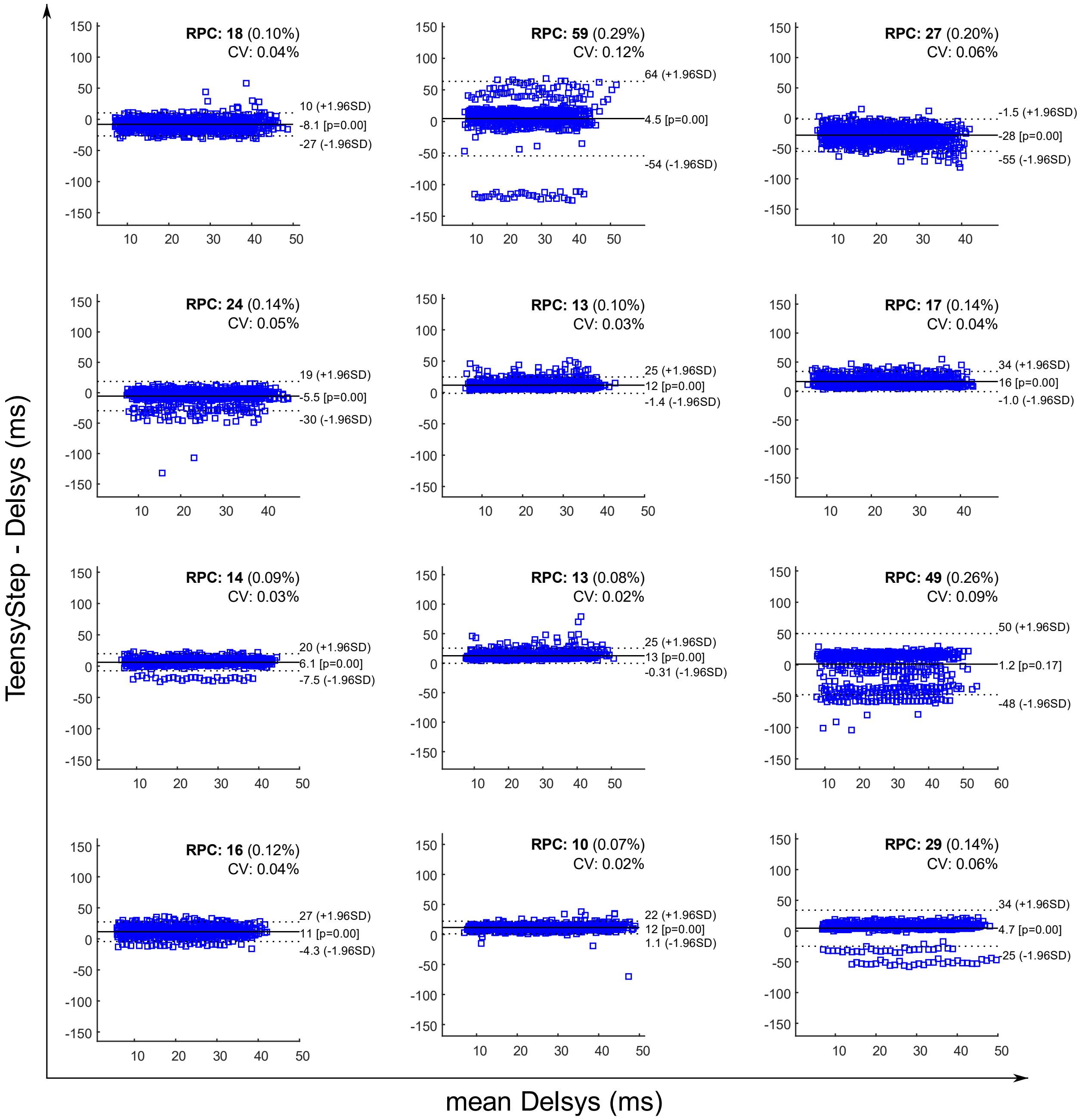
